# Structural Insights on Fusion Mechanisms of Extracellular Vesicles with Model Plasma Membrane

**DOI:** 10.1101/2020.05.25.110601

**Authors:** Fabio Perissinotto, Valeria Rondelli, Beatrice Senigagliesi, Paola Brocca, László Almásy, Laszlo Bottyan, Dániel Géza Merkel, Heinz Amenitsch, Barbara Sartori, Karin Pachler, Magdalena Mayr, Mario Gimona, Eva Rohde, Loredana Casalis, Pietro Parisse

## Abstract

Extracellular vesicles (EVs) represent a potent intercellular communication system. Within a lipid bilayer such small vesicles transport biomolecules between cells and throughout the body, strongly influencing the fate of recipient cells. Due to their specific biological functions they have been proposed as biomarkers for various diseases and as optimal candidates for therapeutic applications. Despite of their extreme biological relevance, the small size (30 to a few hundred nanometers in diameter) of EVs still poses a great challenge for their isolation, quantification and biophysical/biochemical characterization, therefore the complex network of EVs and cells as well as their interaction remains to be further revealed. Here we propose a multiscale platform based on Atomic Force Microscopy, Small Angle X-ray Scattering, Small Angle Neutron Scattering and Neutron Reflectometry to reveal structure-function correlations of purified EVs through the analysis of their interaction with model membrane systems, in form of both supported lipid bilayers and suspended unilamellar vesicles of variably complex composition. The analysis reveals a strong interaction of EVs with the model membranes and preferentially with liquid ordered raft-like lipid domains, and opens the way to understand uptake mechanisms in different vesicle to cell membrane relative compositions.

## Introduction

Extracellular vesicles are nanosized, cell-derived lipid containers devoted to the transport of macromolecules, metabolites and nutrients throughout the body. In the last 15 years they received increasing attention due to their fundamental role in intercellular communication. ^[1]^ EVs are ubiquitously involved in most physiologically relevant processes. Notably, they contain specific signatures from the originating cells and can strongly influence the fate of the recipient cells, hence EVs have been proposed as biomarkers in several diseases. ^[2-5]^ Moreover, their natural biocompatibility, their biological function and their small size make them optimal candidates as therapeutic agents in several frameworks ranging from immune therapy to vaccination, from regenerative medicine to drug delivery. ^[6-7]^ Still, despite their recognized biomedical relevance the field is not yet fully mature and more in-depth studies are required to understand EV physiology. In particular the correlation of biophysical and biochemical properties of isolated EV subpopulations with their biological function is under continuous debate while an overall understanding of EVs cell internalization mechanisms is still lacking. ^[8-11]^ The nanoscale spatio-temporal details on how EVs interact, adsorb, and fuse with target cells, as well as the factors influencing the biogenesis and release of the molecular cargo, are not completely understood. Literature suggests a wide variety of routes for cellular uptake,^[10-13]^ depending on the specific composition of the cellular membrane, EVs function(s) and their physico-chemical properties.^[14,15]^ It is expected that uptake dynamics and membrane fusion mechanisms are tightly related to the potency and function of EVs, and are found to play a key-role in EV-based drug delivery applications.^[16]^ Uptake dynamics in turn has been shown to depend on EV size ^[17]^ and on the extracellular matrix environment,^[18]^ but results on fixed and live cells are quite scattered.

In order to elucidate the EVs/recipient cell interactions here we exploit artificial lipid membranes as tunable model platforms to mimic natural cell membranes. ^[19-22]^ In particular, we challenge our experiments to quantify the dynamics of interaction between fully characterized EVs and model membranes to reveal the relationship between function and biophysical properties of these vesicles. Standardized protocols and Good Manufacturing Practice conditions were employed to derive highly stable vesicles of defined size and reproducible molecular profiles from Umbilical Cord multipotent Mesenchymal Stem (Stromal) Cells (MSCs). After a thorough biophysical and biochemical characterization of EVs non-contact liquid imaging Atomic Force Microscopy (AFM) and, in parallel, Neutron Reflectometry (NR), as well as Small Angle Neutron Scattering (SANS) experiments were performed on EVs to determine their interaction with supported lipid bilayers. As a start, we focused on synthetic membranes constituted by single phospholipids, to further proceed with more complex, 3 components (phospholipids, sphingolipids and cholesterol) systems. The proposed experimental platform provides crucial information on the mechanisms of EVs-cell membrane interaction, as the partial fusion of the EVs with the model membrane bilayer, and on the role played by different lipid phases, paving the way for identifying specific vesicle-cell uptake routes and for modifying them for therapeutic needs.

## Results and discussion

In order to obtain reliable results, we first addressed the EVs isolation for efficiency and reproducibility. A plethora of isolation protocols has been reported, with pros and cons depending on the specific system in use. ^[23]^ In **Figure 1** we present a multi-technique characterization of EVs isolated from Umbilical Cord MSC conditioned medium. Nanoparticle tracking analysis (NTA) shows that EV preparations contain particles with a mean size of 120±5 nm (**Figure 1a**). As confirmed by the clear visualisation of vesicle membrane in cryo-EM images (**Figure 1 c**), these preparations include vesicles. This finding is further supported by the presence of typical EV/exosome markers, such as tetraspanins CD9, CD63 and CD81 in the Multiplex bead-based flow cytometry assay ^[24]^ profiles (**Figure 1 b**). ^[25]^

**Figure 1.**
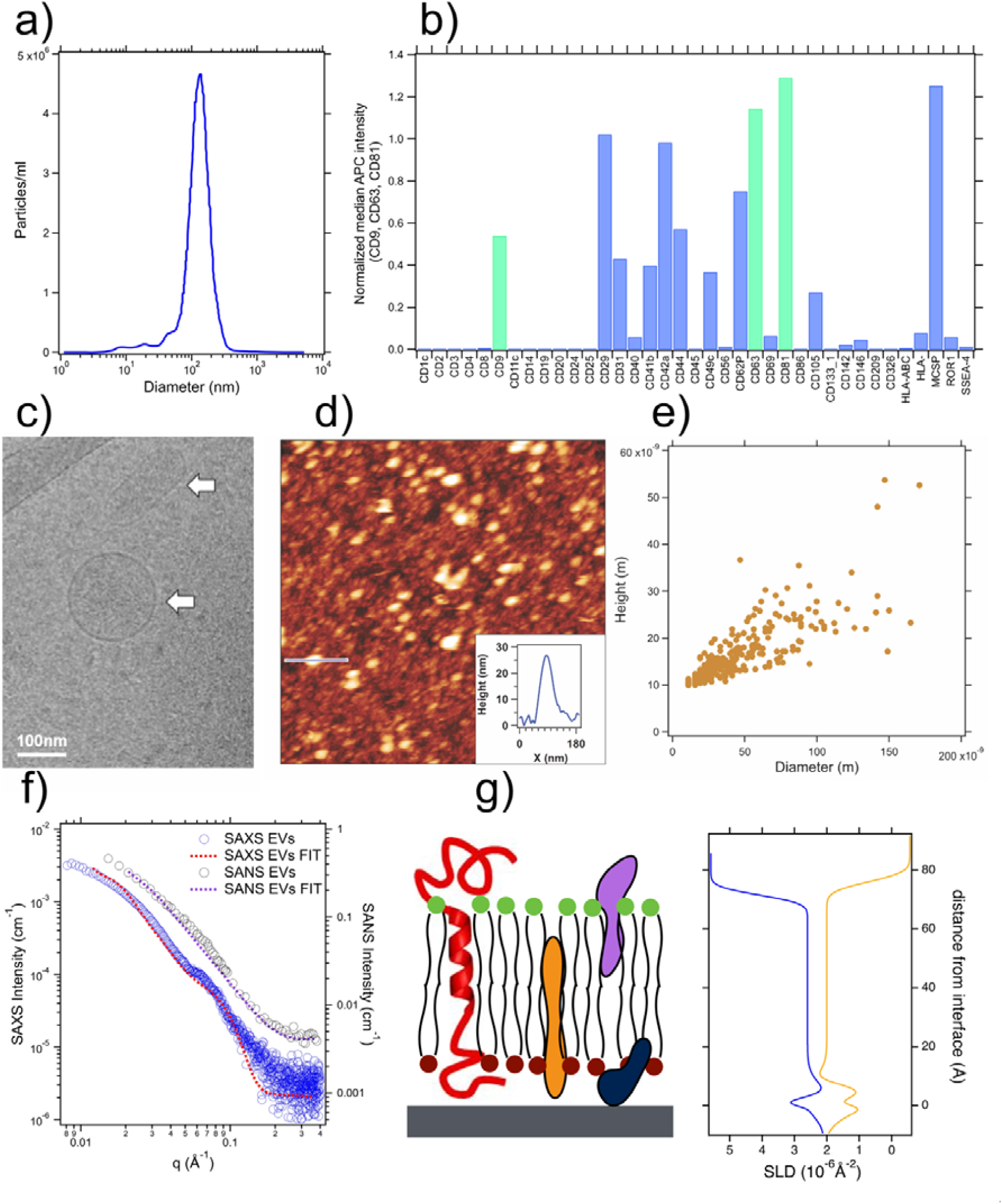
Characterization of isolated Extracellular Vesicles. (a) Size distribution of MSC-derived EVs by nanoparticle tracking analysis. (b). Surface marker profiling by MACSPlex. Standard EV markers CD9, CD63 and CD81 are marked in green. (c) Cryo electron microscopy image from a representative batch of MSC-derived EVs isolated as described in the Methods section. The lipid bilayer surrounding the EV can be unambiguously recognized (arrows) (d) AFM topographic image and corresponding line profile (inset) of MSC-EVs deposited on mica surface. Image acquired in PBS. (e) Correlation diagram between the diameter and the height extracted from AFM analysis. (f) SANS (black circles) and SAXS (blue circles) profiles of MSC-derived EVs and corresponding fits (dark pink dotted line and red dotted line) with a core-shell model. (g) Scattering Length Density profiles of a EVs-derived SLB in H_2_O (orange) and D_2_O (blue) obtained by Neutron Reflectivity with a pictorial sketch to help in SLDs profiles interpretation. Measured reflectivity data and best fit parameters are reported in the SI.

AFM profiles from topographic images in liquid (**Figure 1 d, e**) confirmed the size distribution of EVs observed by NTA and cryo-EM analysis, highlighting the presence of vesicles sized less than 50 nm, not detectable by NTA. In **Figure 1 e** the vesicles are displayed to exhibit diameters ranging between 30 and 150 nm and heights ranging from 10 to 40 nm. The slightly deformation with respect to their supposed spherical shape may be attributed to the small force applied by the AFM tip, the immobilization on the mica surface and possible tip convolution effects. ^[26-28]^

To gain further information on the structural properties of EVs we performed Small Angle X-ray Scattering (SAXS) and SANS experiments on EVs in solution (**Figure 1 f**). Analogous approaches have been applied to investigate similar systems. ^[29-31]^ The SAXS profile in the investigated q-range, analysed by a simple core multi-shell model, is consistent with an asymmetric membrane profile, reported in **Figure S1**, accounting for extended polar components. However, the detailed feature at q values around 0.06-0.08 Å-1 may originate from some characteristic distance in the range of 10 nm occurring among objects on membrane surface, that can agree with the average pattern of rugosity observed by AFM. The same pattern may determine the detected SANS intensity, in the analogous q-vector range, deviating from the profile of a shell of about 2 nm thickness (fitting curve in **Figure 1**) representing the scattering from the membrane core.^]^

To further characterize EVs we performed NR studies. To prepare NR samples, EVs were deposited on the surface of a macroscopic silicon support, on which they fused. The EVs-derived membrane reached a total surface coverage of 90%. A layer of 0.5 nm water was measured between the membrane and the silicon support. The cross structure of the supported bilayer was probed by NR, in analogy with what has already been performed on model membranes as well as bacteria extract systems. ^[32,33]^ **Figure S2 a** displays NR scan and best fit, while in **Figure 1 g** we report the corresponding scattering length density (SLD) profiles. NR data on EVs with two different solvent contrasts were collected and simultaneously fitted (**Figure 1** g, D_2_O, blue line; H_2_O yellow line). Data analysis confirmed complete fusion of EVs on the silicon surface giving rise to a membrane of 6.9 ± 0.2 nm thickness, with a SLD of 2.5 ± 0.2×10^−6^ Å^-2^. The SLD value is consistent with a mixture of lipids (SLD ≈ -0.5 − 1 x10^−6^ Å^-2^) and proteins (SLD ≈ 1.5 − 3 x 10^−6^ Å^-2^). Although higher than the thickness of both synthetic and natural lipid extracts which is generally around 5.5 nm, ^[34-36]^ the thickness of the EV-based bilayer is compatible with one single bilayer. Further, the scattering profile is consistent with the presence of molecules other than lipids, as large proteins, as part of the deposited membrane. The system interfacial roughness was found as low as 0.6 ± 0.2 nm. This finding itself validates the entire approach, giving robustness to the data analysis model and confirming a successful deposition of EVs.

In order to investigate the interaction of EVs with cell membranes, we used planar supported phospholipid bilayers as model membranes that we characterized by thorough AFM topographic imaging. In **Figure 2a** a representative AFM image and the corresponding line profile of a 1,2-dioleoyl-sn-glycero-3-phosphocholine (DOPC) supported lipid bilayer (SLB) are reported. DOPC was chosen for its extreme homogeneity, for its almost defect-free structure, and its very small surface roughness (0.17±0.05 nm). The bilayer height, as measured from indentation profiles and topographic measurements of partially complete bilayers (see **Figure S3**), is 5 ±0.2 nm in agreement with literature reports. ^[37]^

**Figure 2.**
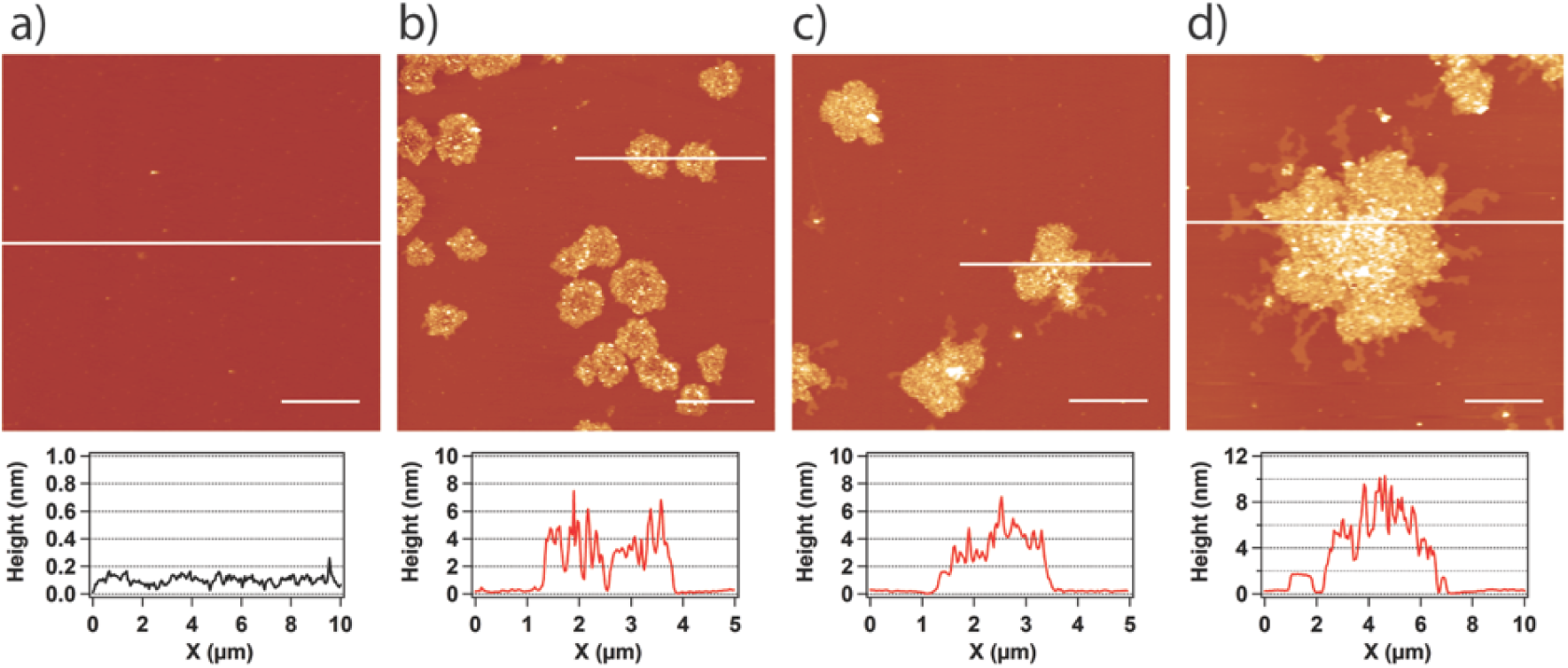
((AFM topographic images of a DOPC-SLB before addition of EVs (a) and past 30 min (b), 90 min (c), and 210 min (d) exposure times following EV addition. The cross-sectional line profile of (a) proves the high homogeneity and flatness of the DOPC SLB and those of (b,c,d) the protrusion of EVs islands inside the DOPC SLB. Lateral scale bar is 2 μm.)

The DOPC SLB topographic image in **Figure 2 a** is then taken as the time zero snapshot of a time resolved AFM experiment to monitor the bilayer modification upon EV addition. After 30 minutes following the addition of EVs one clearly observes the presence of islands protruding 3-4 nm from the DOPC surface. On the one hand, there was no trace of intact EVs. Note that the ∼120 nm diameter of the EVs would be sufficient to make them recognized by topographic imaging, as shown in **Figure 1 d** for EVs on mica. On the other hand, the observed protrusion of the islands is less pronounced – namely 3.4±0.7 nm vs. 5.5±0.5 nm – than what might be expected if a second lipid bilayer starts to develop on top of the DOPC bilayer.. Therefore, we interpret this mesoscale heterogeneity to be a consequence of presence of membrane proteins as well as the cargo macromolecules released from the opened EVs in the modified but still single membrane bilayer. ^[38]^ A similar mesoscale heterogeneity has been observed also in silica-supported EV-derived SLB, ^[39-40]^ and has been interpreted with the distribution of different portions of the cargo trapped in the SLB.

As a function of EV exposure time (**Figure 2 c,d**) the islands grow in lateral dimensions (from a few hundred of nanometers to micrometer diameter), until they coalesce into larger islands (of about 3-4 micrometer diameter after 4 hours of exposure time). The growth evolution of the EV-enriched patches in the AFM images is in agreement with that of the formation of EV-derived SLB, as reported by Montis and coworkers. ^[39]^

The single bilayer hypothesis for the EV-related islands implies a (partial) mixing of EVs with SLB and it might be considered a preliminary step towards complete fusion of the EVs with the membrane itself. Lipid mixing is also supported by the appearance of a new lipid phase around the EV patches, which protrude by about 1 nm (Figure 2 d), after 4 hours exposure time. Such a phase might be assigned to the partial diffusion of EVs’ lipids in the DOPC layer and/or to a misalignment of DOPC molecules due to the presence of proteins or other molecules diffusing under the bilayer. A similar mechanism of membrane fusion has been also hypothesized, based on combined atomic force and fluorescence microscopy experiments, for proteoliposomes interacting with SBL, ^[41]^ in accordance with our findings. Yet, a full understanding of the fusion mechanism would require to discriminate possible fusion asymmetries occurring in the two membrane leaflets. In order to prove or deny the presence of proteins in the EV-related lipid phase, following a proteinase K treatment using the protocol of Skliar *et al.*, ^[42]^ we incubated the treated EV to a DOPC SLB and monitored the morphology of the system for the same duration as indicated in **Figure 2 (b)**. From AFM topographic imaging (**Figure S4)** we observed the presence of patches with lateral dimensions comparable to those originated by non-treated EVs. However, the average height is now roughly the same of the DOPC layer, although with a sensibly increased roughness. Altogether these findings indicate that i) the EVs completely open on the SLB; ii) the topographical heterogeneity observed in Figure 2, is attributable to EV proteins; iii) upon addition of EVs a mixed SLB forms in which the lipid components of the EV tend to remain segregated /intercalated in the DOPC SLB.

To further clarify the interaction of EVs with the PC lipid bilayer, and investigate the extent of fusion, that is if fusion involved only the external target membrane leaflet or the whole membrane, we performed NR and SANS measurements. NR allowed to investigate the transverse structure of a single supported deuterated phospholipid (d54-DMPC) bilayer before and after the interaction with EVs. The phospholipid membrane was prepared by vesicle fusion. The d54-DMPC was deuterated in order to exploit the proton-deuteron (actually hydrogen-deuterium, H-D) contrast difference in neutron scattering. This approach was previously adopted by our group when monitoring the interaction of macromolecules, as alpha-synuclein, with artificial bilayers, ^[43-45]^ similar to the method used by Ghosh *et al*. to study synaptic vesicles. ^[46]^ First the neutron reflectivity of the bulk deuterated bilayer was recorded, then, a solution containing isolated EVs was added to the cell with the same EV/lipids ratio used in the AFM experiments and the NR was recorded again. By comparing the reflectivity profiles, one expects to see how the H-rich EVs penetrate the deposited D-rich bilayer. The measured reflectivity profiles together with the simultaneous best-fits are reported in **Figure S2**, while the relative SLD profiles are shown in **Figure 3**.

**Figure 3.**
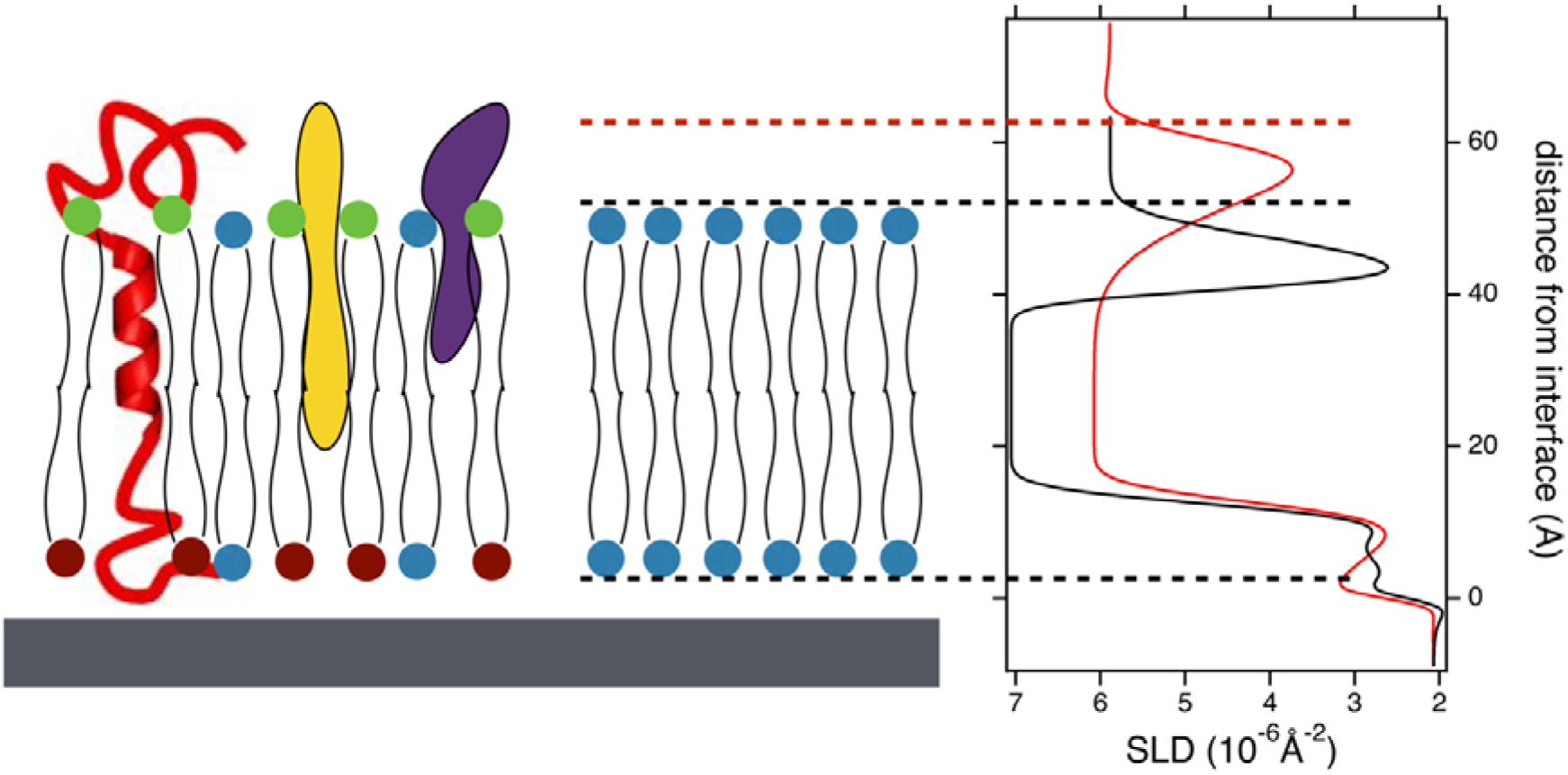
Scattering Length Density profiles of a deuterated PC SLB before (black) and after EVs interaction (red) obtained by Neutron Reflectivity with a pictorial sketch to help in SLDs profiles interpretation. Measured reflectivity profiles and best fit parameters are reported in the SI.

The analysis of the obtained SLD profiles shows an overall increase of the thickness of the membrane upon EV-addition and an overall modification of the SLD profile across the membrane. The increased membrane thickness of 5.4±0.1 nm (from 4.1±0.1 nm in the bulk bilayer) is an intermediate value between that of a pure phospholipid membrane and of an EVs-based supported membrane (6.9 nm), as reported in **Figure 1 g**. We also observed a decrease of SLD value of the membrane from 7.1±0.2 x 10^−6^ Å^-2^ to 6.1±0.2 x 10^−6^ Å^-2^, consistent with the fusion of the H-containing EVs into the deuterated membrane with 20% volume penetration. This experiment demonstrates the high fusogenic ability of EVs with the lipid-only target membrane. Moreover, the fusion occurs “transmembrane”, i.e. in the entire thickness of the bilayer, involving not only the external, but also the inner layer (with respect to the solid support) of the bilayer. Interestingly, the final membrane profile becomes slightly asymmetric (see **Figure 3**). This asymmetry may reflect an actual uneven or rough feature of EV fusion with the bilayer. We cannot exclude at this point, however, a contribution from the silicon support, generally negatively charged in aqueous environment, thus impeding the extended hydrophilic molecular portions to position themselves in the inner leaflet.

To deepen this aspect, we performed fusion experiments in bulk applying SANS to study the structural details of mixed solutions containing unilamellar phospholipid vesicles and EVs in different proportions (details in the materials and methods section). Again, deuterated phospholipid vesicles were used to benefit the H-D contrast difference and to reduce the incoherent background due to the H atoms. **Figure 4** a) summarizes the SANS results on the low, medium and high concentration vesicle-to-EV mixtures of number ratios 15000:1, panel (a); 3000:1, panel (b) and 2700:1, panel (c), respectively. The measured intensities can be compared with those of the pure phospholipid vesicles (red circles) and pure EVs (black circles) as well as their weighed sum in the respective proportions (dotted lines). Panel (c) displays the SLD distributions derived from the fit of the SANS data. Despite the low molar proportion of EVs with respect to target vesicles, as visible from experimental results, the H-bringing EVs scattering intensity is highlighted within the D-based phospholipid target membranes, allowing for a detailed structural investigation of the mixed systems.

**Figure 4.**
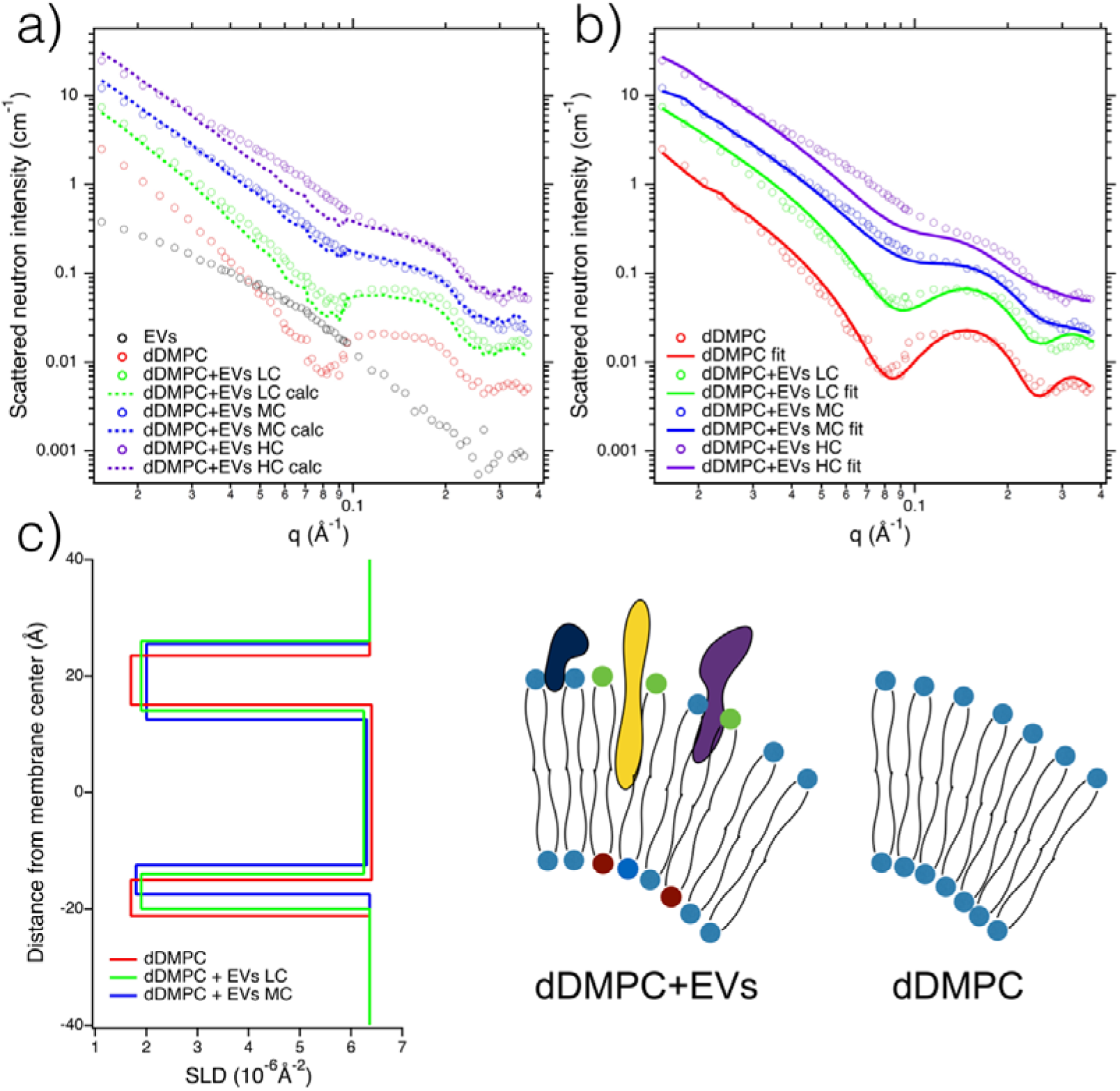
SANS measurements on deuterated PC unilamellar vesicles at 20 mg/mL (red circles), EVs at the concentration of 2*10^9^ EVs in 100 µL (black circles) and the mixed systems at low (green circles), medium (blue circles) and high (violet circles) relative proportions, see text. a) Dotted lines represent the weighed sum of the spectra of the two pure components (phospholipid vesicles and EVs) of appropriate proportions. b) Continuous lines are the fits of the experimental curves with a three-layered vesicular model, parameters in SI. c) SLD profiles extracted from fits to the SANS data (see SI).

In **Figure 4 a** the shape of the reconstructed and of the measured curves are different, indicating a non-negligible interaction between the constituents. For medium and high EVs concentration, the experimental scattering profiles cross the reconstructed ones, indicating that a change in the SLD of the objects occurred. This suggests a decrease of the overall size of the mixed systems with respect to the original vesicles, as shown by the lower intensity at low q vectors.

Hypothesizing that the interaction leads to a unique mixed vesicle distribution, we tentatively analysed the data by modelling the system with the form factor of a (three) layered vesicular system (outer hydrophilic, hydrophobic and inner hydrophilic layers) with a water core. This simplified model seems to work well for systems with null or low EV content and works increasingly less for systems with larger EVs concentration. Best fit results and parameters are reported in **Figure 4b** and **table S3**, while the SLD profiles of the single bilayer obtained by the data best fits are shown in **Figure 4 c**. Data analysis is consistent with a lowering of the SLD of the hydrophobic portion, indicating that the mixing involves both membrane leaflets, in accordance with the NR results. Moreover, the thickness of the hydrophobic portion appears to decrease upon EVs mixing, while the thickness of the external polar portion increases at increasing EVs content. Results on thicknesses alteration agree with both AFM and NR findings. Furthermore, the analysis is consistent with the formation of an asymmetric membrane upon EVs mixing. The external hydrophilic layer becomes more extended after fusion, as if the largest membrane components belonging to EVs were not included in the inner membrane leaflet of the final vesicular system, but only fuse to the outer one, as already observed by NR and depicted in **Figure 3 c**.

These findings unveil that the mixing mechanism is not a total fusion, rather lipids and smaller molecules might fuse and eventually flip into the inner membrane leaflets, while large proteins bringing important hydrophilic portions could eventually reside in the outer membrane leaflet. Notably, this asymmetry is kept stable for a long time (2 days of measurements), suggesting that the flip-flop is not energetically favoured. When considering this limit to total fusion it is important to recall that so far we have been dealing with simple models of target membranes. Rather, in natural biological membranes, the lateral complexity related to the occurrence of ordered domains of different lipids and to the presence of membrane proteins plays a significant role in fusion mechanisms. Ascertained the intercalation and breaking of EVs inside the liquid disordered phase of the PC bilayer, to address real membrane complexity we prepared a mixed bilayer composed of DOPC:Sphingomyelin (SM) 2:1 with a 5% cholesterol, mimicking the lipid ratios of neuronal membranes. According to the relative three-component phase diagram, such mixed SLB presents, at room temperature, a phase separation between domains in liquid ordered (*Lo*) phase enriched in SM and domains in liquid disordered (*Ld*) phase enriched in PC. ^[47]^

AFM analysis reported in **Figure 5 a**, shows the presence of domains protruding 1 nm above the surrounding bilayer, in accordance with literature reports on the expected difference in height between *Lo* and *Ld* domains. ^[47, 48]^ The mixed lipid surface is slightly more defective than the pure DOPC bilayer, also in line with the literature. We then exposed the SLB to EVs, using the same experimental conditions used for the DOPC SLB. In **Figure 5 b-c** we report the temporal evolution of the interaction as measured by AFM. Even in this case we can observe the formation of patches on the surface that tend to coalesce in larger domains as time passes, with the EV-related domains height roughly protruding 4 nm above the SLB. Interestingly the EV-related domains seem to co-localize with the SM enriched domains. The analysis of the areas covered by SM before and after EV interaction points to an almost complete disappearance of the *Lo* domains at advantage of the EV patches.

**Figure 5.**
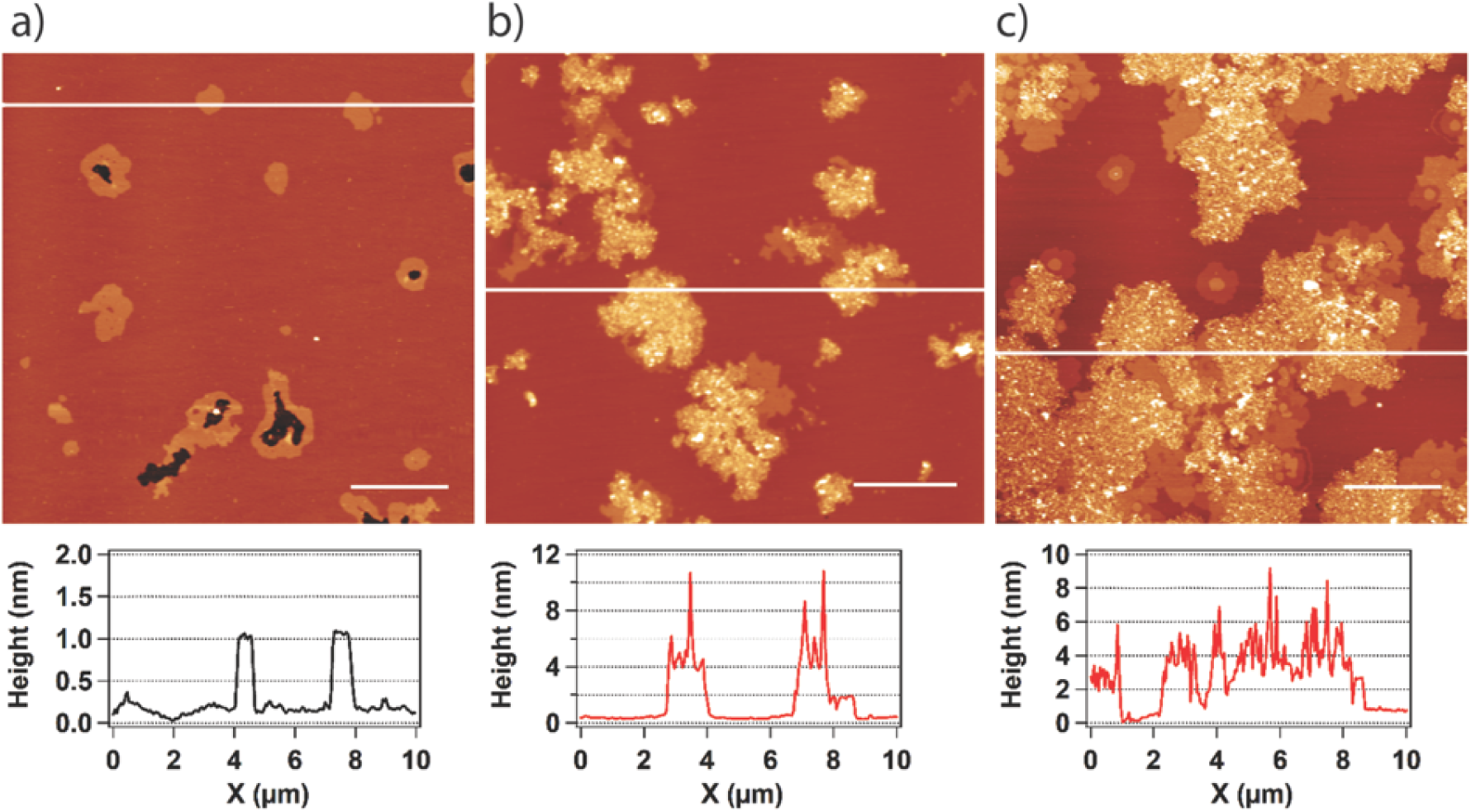
AFM topographic images of a DOPC:SM:Chol (2:1:0.15) SLB before (a) and after addition of EVs (b,c). We report the temporal evolution of the formation of EVs SLB inside the DOPC SLB after 30 min (b), and 240 min (c). We also report the cross-sectional line profiles to highlight the high homogeneity and flatness of the artificial SLB with the 1 nm difference in height between liquid disordered and liquid ordered domains (a) and the protrusion of EVs SLB inside the artificial SLB in (b-c).)

The observed preferential interaction of EVs with SM-enriched domains is compatible with the biological mechanism of lipid-raft mediated endocytosis and/or with the preferential docking of EVs in regions of the SLB where a discontinuity is present. ^[49,50]^. Moreover, comparing the temporal evolution of the coverage of EVs SLB inside the artificial bilayers (pure DOPC and DOPC:SM:Chol) we observe that both follow a first-order Langmuir absorption behaviour (**Figure S4**), with similar kinetic constant but with a larger final coverage for the mixed bilayer, indicating an higher affinity of the EVs towards the raft-like membrane.

## Conclusions

Here, we provided for the first time a molecular description of the interaction of EVs with SLBs as model plasma membrane systems. We showed that AFM morphological analysis can be profitably combined with neutron scattering-based structural investigation to successfully elucidate the molecular mechanisms of interaction of EVs with membranes, from which EVs cargo release may depend. In particular we demonstrated the EVs propensity to fuse with the model membranes with a preferential interaction with the external layer of the fluid membrane. Also, we highlighted the critical role of ordered lipid compartments mimicking lipid rafts to promote EVs fusion. The use of variable composition SLBs as model systems to mimic cell membranes will be particularly useful in studies of EVs-cell uptake, in which concurrent multiple uptake channels as lipid-raft mediated endocytosis, surface binding and membrane fusion might be involved ^[12]^, allowing to disentangle the various contributions addressing them one at the time. These different mechanisms are in fact particularly difficult to be distinguished in a complex cellular environment. ^[51-53]^ Knowledge acquired from model systems on the prevalent uptake mechanism of distinct EV subpopulations and on the specific membrane subdomain involved will allow to design specific strategies to act on the defined internalization channel and/or membrane compartment to block or favour cell EV interactions and cargo release to target cells in the context of a given disease. The approach shown here can be extended to convey incremental complexity, adding glycolipid and membrane proteins to the model lipid bilayers. We strongly believe that our approach combined with data on the specific biological function of each EV subpopulation as retrieved by standard functional assays, will turn useful to select the crucial molecular aspects of EVs internalization by cells. Such information might be then exploited to develop novel devices based on standard readout platforms, e.g. fluorescence, mechanical and/or electrochemical readout, to screen EVs for particular properties to predict their functionality, thus revolutionizing the way how EVs may be used in diagnostics and hopefully become useful in therapy.

## Experimental Section

### Manufacturing of human mesenchymal stromal cell–derived extracellular vesicles (EVs)

Human umbilical cord-derived mesenchymal stromal cells (UC-MSCs) were seeded (1560 cells / cm^2^) into a CF4 cell factory system (Nunc) in fibrinogen-depleted culture medium composed of α-MEM (Sigma-Aldrich), 10 % pHPL, and 5 mM (N2)-L-alanyl-L-glutamine (Fresenius Kabi)^1–3^. Cells were cultured at 5 % CO_2_ at 37 °C. Upon reaching a cell confluence of 60 – 70 %, cells were washed with phosphate-buffered saline (PBS) solution and the growth medium was exchanged to a EV-harvest medium, consisting of α-MEM, 5 % pHPL – (para-hydroxyphenyllactate), which was additionally EV-depleted by tangential flow filtration (TFF) using a 750 kDa hollow fibre filter (Spectrum Labs), and 5 mM (N2)-L-alanyl-L-glutamine. After 24h, conditioned medium was harvested, centrifuged at 2.500 x g for 20 mins at 18 °C, and filtered (0.22 µm). EVs were concentrated and buffer-exchanged to PBS by TFF using a 100 kDa hollow fibre filter (Spectrum Labs). Ultimately, EVs were isolated and concentrated by ultracentrifugation at 120.000 x g for 3 h at 18 °C. Resulting EV-pellets were washed with PBS and subsequently resuspended in Ringer’s Lactate in an appropriate volume to achieve a dose of 20 – 40 x 10^7^ / mL (cell equivalent). Resuspended EVs were centrifuged at 3.000 x g for 10 min at 4 °C, sterile filtered (0.22 µm), and stored in glass vials at – 80 °C.

In certain cases, EVs were further purified by size exclusion chromatography (SEC) using qEV original SEC columns (Izon Science) according to the manufacturer’s instructions.

### Lipids for model membranes

1,2-dimyristoyl-sn-glycero-3-phosphocholine (DMPC), 1,2-dimyristoyl-d54-sn-glycero-3-phosphocholine (d54-DMPC), 1,2-dioleoyl-sn-glycero-3-phosphocholine (DOPC), sphingomyelin (brain, porcine, SM) and cholesterol (ovine wool, > 98%) have been purchased by Avanti Polar Lipids (Alabama) and used without any further purification.

### Nanoparticle Tracking Analysis

For nanoparticle tracking analysis of EV samples in light scatter mode, EVs were diluted to a concentration of 4 – 7 x 10^7^ particles / mL in PBS. Prior to EV-sample analysis, the instruments were calibrated using YG-labeled 100 nm polystyrene standard beads (1:1.000.000 dilution in ddH_2_O). For calibration of the PMX 110 instrument, minimum brightness was appointed to 25 AU (arbitrary units), temperature to 21.5 °C, shutter to 70 AU, and sensitivity to 65 AU. The minimum brightness of the PMX 120 instrument was set to 30 AU, temperature to 21.5 °C, shutter to 100 AU, and sensitivity to 65 AU.

To determine the size and number of particles in the prepared EV-samples, EV samples (fluorescently-labeled or unlabeled) were analyzed in light scatter mode. For the PMX 110 instrument, the minimum brightness was therefore set to 20 AU, temperature to 21.5 °C, shutter to 70 AU, and sensitivity to 85 AU. Subsequently, data for two exposures at 11 measurement positions were acquired per sample. Afterwards, data for two exposures at 11 measurement positions were collected for each sample. Based on the Stokes-Einstein equation, particle sizes were determined using the ZetaView software (PMX 110: Version 8.4.2, PMX 120: Version 8.5.5).

### Macs Plex

The MACSPlex Exosome Kit (Miltenyi) is a bead-based multiplexed FACs based assay for the analysis surface markers present on EVs. We have used the MACSPlex kit according to the manufacturer’s instruction and following a validated standard operating procedure with 5 x 10^7^ to 5×10^8^ particles as input. Data acquisition was conducted on a FACS Canto II (BD Biosciences).

### Unilamellar Lipid Vesicles Preparation

Model lipid-based membranes were prepared according to a standard procedure explained elsewhere ^[54]^ by thin film deposition, hydration and extrusion through policarbonate filters with controlled porosity of 80 nm.

### Atomic Force Microscopy

AFM imaging was carried out on a commercially available microscope (MFP-3D Stand Alone AFM from Asylum Research, Santa Barbara, CA) working at room temperature in dynamic AC-mode. Commercially silicon cantilevers (BL-AC40TS-C2, Olympus Micro Cantilevers, nominal spring k 0.09 nN/nm) have been chosen for imaging in liquid. Images were acquired at 512 x 512 pixel frames at 0.6-1.0 lines/s scan speed.

For AFM analysis of EVs molecules, 10 µL of EVs were spread onto 0.6 cm x 0.6 cm piece of freshly cleaved mica, left to incubate for 5 min, gently rinsed 3 times with PBS, and imaged by AFM in liquid.

For EVs membrane-interaction investigation we prepared two different model membranes: one made by DOPC phospholipid and one made by DOPC+SM+Chol in 2:1:0.16 molar ratio. 100 µL of SUVs solution (0.5 mg/mL in 2 mM CaCl_2_) of specific lipid composition were spotted on freshly cleaved mica attached to the AFM liquid chamber by ultrafast glue. The sample was left to incubate for 15 minutes at room temperature to promote vesicle adsorption, fusion and the formation of lipid bilayer on the surface. Then, lipid membrane was gently rinsed three times with Milli-Q H_2_O to remove the excess of vesicles from the liquid sub-phase before AFM analysis.

EVs have been inserted in the AFM liquid cell and the measurements were acquired 15 minutes after incubation. AFM image analyses were performed using Gwyddion, an open-source modular program for scanning probe microscopy data visualization and analysis ^[55]^. Graphs representing AFM trace profiles and height distributions were obtained using Igor Pro software (Wavemetrics, US).

### Small Angle X-Ray Scattering

Small Angle X-Ray Scattering (SAXS) experiments have been performed at the Austrian SAXS beamline in Elettra Synchrotron in Basovizza, IT. ^[56]^ The sample to detector distance was set to 1200 mm, allowing to investigate a q-range between 0.08 and 5.5 nm^-1^. MSC-EVs were resuspended in phosphate buffer solution (PBS), to the final concentration of 2*10^9^ EVs in 100 µL, inserted in a quartz capillary and measured at 25°C.

### Neutron Reflectometry

d54-DMPC has been used to prepare model membranes to perform Neutron Reflectometry (NR) experiments. The use of deuterated phospholipids helps in enhancing the visibility of molecules interacting with the lipid bilayer, thanks to the induced membrane contrast change, being the scattering length density of the deuterated lipids around 7×10^−6^Å^-2^, while that natural lipids around -0.5×10^−6^Å^-2^ and that of proteins around 2.5×10^−6^Å^-2^.

NR measurements were carried out at the GINA neutron reflectometer at the Budapest Neutron Centre (BNC). ^[57,58]^ The solid-liquid interface was investigated at a neutron wavelength of 4.63 Å in an aluminum liquid cell (temperature controlled by water circulation to ±0.1°C) which held a 70×70×10 mm^3^ Si (100) block with a smooth surface (rms roughness of 0.3 nm) for membrane fusion and investigation. The neutron beam entered the cell from the Si block side. The single supported d54-DMPC membrane was obtained by the fusion of lipid vesicles injected in the cell at the concentration of 0.5 mg/mL. Prior to the NR experiment, the Si block of the cell was cleaned by organic solvents, then cleaned in a plasma cleaner for 15 minutes and finally washed with Milli-Q H_2_O.

For EV-based bilayer investigation, after support characterization, a solution of 3*10^9^ EVs in 2 mL of D_2_O have been injected in the measuring cell (10 mL total volume) at T = 37 °C and, after 45 min for incubation, the excess material has been gently removed by flushing D_2_O in the cell. Finally, reflectivity has been measured in two contrast solvents (H_2_O and D_2_O).

### Small Angle Neutron Scattering

d54-DMPC has been used to prepare model membranes to perform the SANS experiments. The use of deuterated phospholipids helps in enhancing the visibility of molecules interacting with the lipid bilayer, thanks to the induced membrane contrast change, being the scattering length density of the deuterated lipids around 7×10^−6^Å^-2^, while that natural lipids around - 0.5×10^−6^Å^-2^ and that of proteins around 2.5×10^−6^Å^-2^.

Measurements were performed on the SANS-YS instrument at the Budapest Neutron Centre using two wavelengths and two sample-to-detector distances to cover a wide q-range, from 0.005 to 0.4 Å^-1^. Lipid membranes have been prepared in the form of extruded monolamellar vesicles, while EVs were resuspended in D_2_O (10^10^ vesicles/ml) Samples were placed in quartz cells (produced by Hellma) and measured at 25°C.

For fusion experiments, 500 µL of phospholipid vesicles at the concentration of 20 mg/mL have been mixed to 100 µL of EVs solution (3*10^12^ phospholipid vesicles + 2*10^8^ EVs (LC)); to 500 µL of EVs solution (3*10^12^ phospholipid vesicles+ 10^9^ EVs (MC)) and to 600 µL of EVs solution (3*10^12^ phospholipid vesicles + 1.2*10^9^ EVs (HC)).

Deuterated lipids have been chosen to highlight the presence of eventual H-containing molecules (by which EVs are composed) within the model membranes in D_2_O.

Data fits have been performed by the software SasView. ^[59]^

## Supporting Information

Supporting Information is available.

## Acknowledgements

The authors acknowledge funding from the European Regional Development Fund Interreg V-A Italia–Austria 2014–2020 (EXOTHERA ITAT1036). We acknowledge CERIC-ERIC proposal grant n. 20187082 to perform measurements at the Austrian SAXS beamline of Elettra Sincrotrone Trieste S.C.p.A. and at GINA and SANS-YS instruments of the Budapest Neutron Center. High purity cleaning of the Si block surface by the staff of the BioMEMS group of Inst. Tech. Physics and Mater. Sci., Budapest is gratefully acknowledged. We gratefully acknowledge the assistance of H.M. Binder and D. Auer (PMU Salzburg) for flow cytometry analyses, and A. Desgeorges (PMU Salzburg) for NTA analyses.

## Notes

### Competing Interest Statement

The authors have declared no competing interest.

