## supplementary material for "Structural Insights on Fusion Mechanisms of Extracellular Vesicles with Model Plasma Membrane"

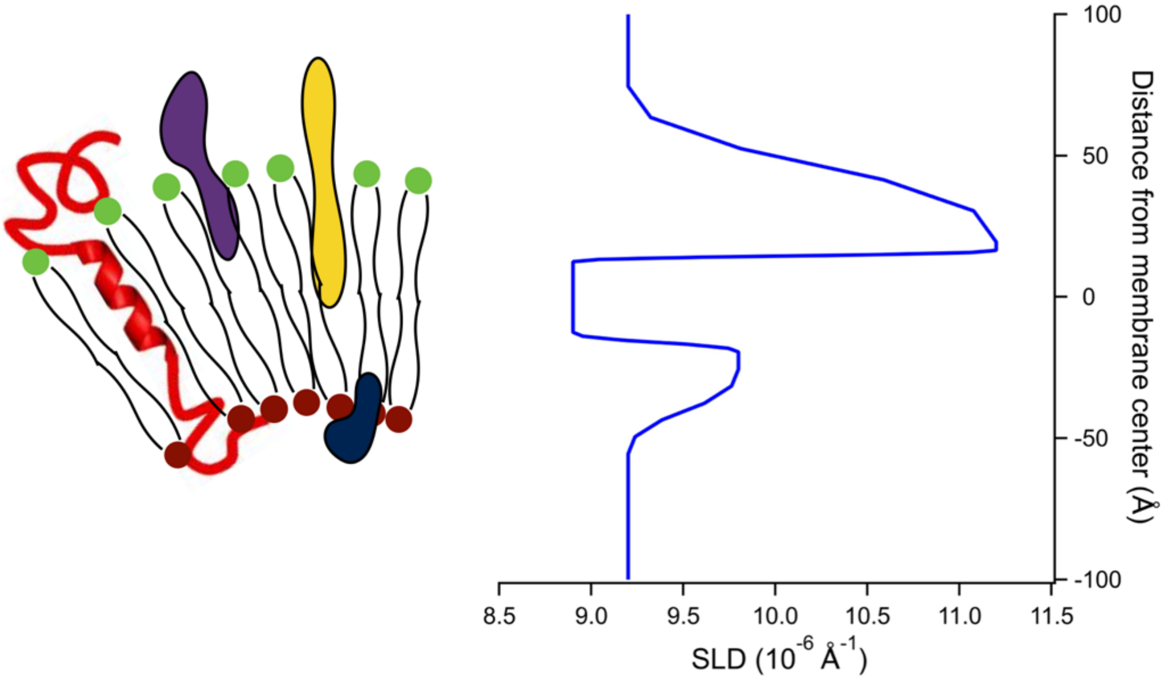


**Figure S1. EVs Scattering Length Density profile** extracted from SAXS data fits obtained with a 3-layered membrane model, accounting for the proteomic component extending in the extra-vesicular solution.

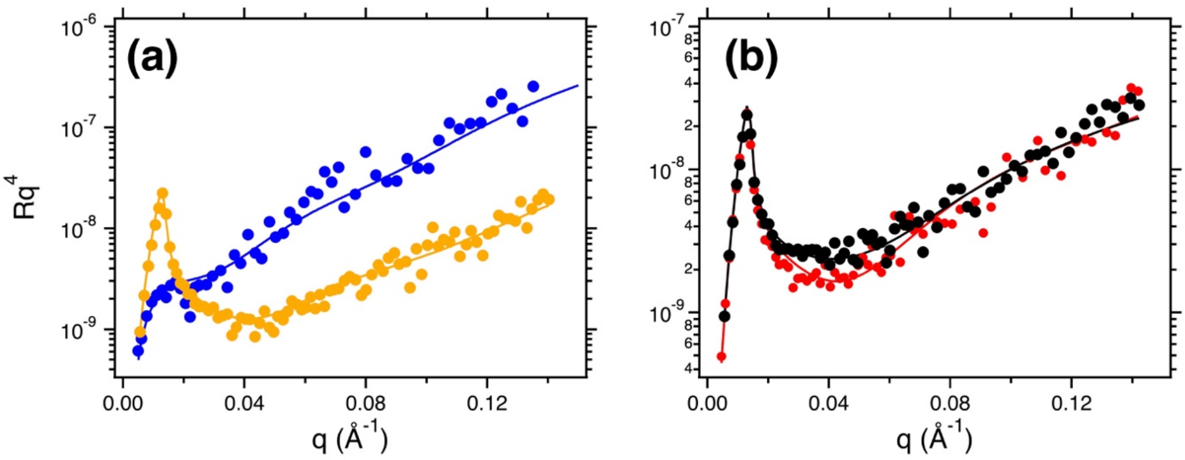


**Figure S2. Neutron reflectometry from EVs based supported bilayers**. a) Experimental data and relative fits (Left panel) of EVs based supported membranes investigated in two contrast solvents: $H_2O$ (blue) and $D_2O$ (orange). The system has been investigated at 37°C. Best fit parameters are reported in Table S1. b) Experimental data and relative fits (Left panel) of d54-DMPC SLB before (black) and upon addition of EVs (red). The system has been investigated at 37°C. Best fit parameters are reported in Table S2.

| | **Thickness (± 2 Å)** | **SLD (± 0.2 $E^{-6}$ $Å^{-2}$)** | **Roughness with previous layer (± 2 Å)** |
|---|---|---|---|
| **$SiO_2$** | 1,5 | 3,4 | 2 |
| **solvent** | 5 | | 6 |
| **membrane** | 69 | 2 | 2 |

**Table S1.** Best fit parameters for NR curves of EVs based supported bilayers

| | **DMPC** | | | |
|---|---|---|---|---|
| | **Thickness (± 2 Å)** | **SLD (± 0.2 $E^{-6}$ $Å^{-2}$)** | **Solvent penetration (± 3% vol)** | **Roughness with previous layer (± 2 Å)** |
| **$SiO_2$** | 1.5 | 3.4 | | 2 |
| **solvent** | 5 | | | 3 |
| **Heads in** | 7 | 1.75 | 15 | 2 |
| **Chains** | 28 | 7.1 | 4 | 2 |
| **Heads out** | 7 | 1.75 | 15 | 3 |

| DMPC + EVs | | | | |
|---|---|---|---|---|
| | Thickness (± 2 Å) | SLD (± 0.2 E$^{-6}$ Å$^{-2}$) | Solvent penetration (± 3% vol) | Roughness with previous layer (± 2 Å) |
| **$SiO_2$** | 1.5 | 3.4 | | 2 |
| **solvent** | 5 | | | 5 |
| **Heads in** | 7 | 1.8 | 15 | 4 |
| **Chains** | 40 | 6.1 | 4 | 2 |
| **Heads out** | 7 | 1.8 | 15 | 7 |

**Table S2.** Best fit parameters for NR curves of DMPC and DMPC + EVs supported bilayers.

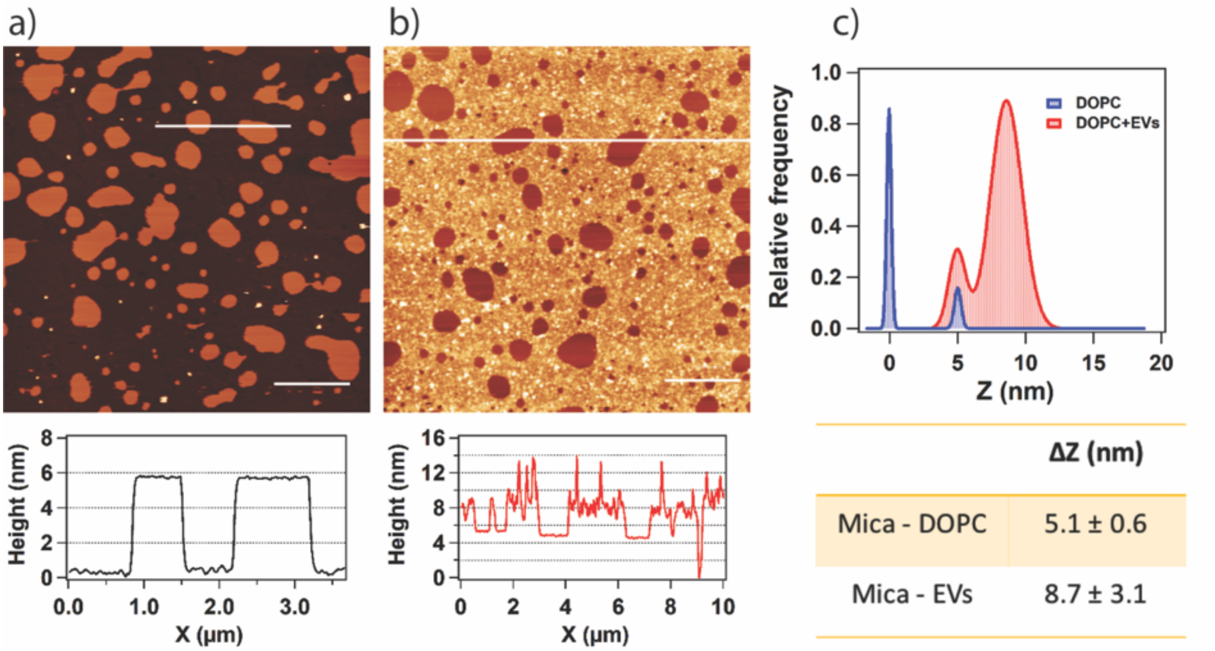


**Figure S3.** Atomic Force Microscopy topographic images and corresponding line profiles of a partial DOPC SLB before (a) and after addition of EVs (b). We report in (c) the height histograms relative to the two AFM images, to highlight the height of the DOPC SLB (a) and the protrusion of EVs SLB inside the DOPC SLB.

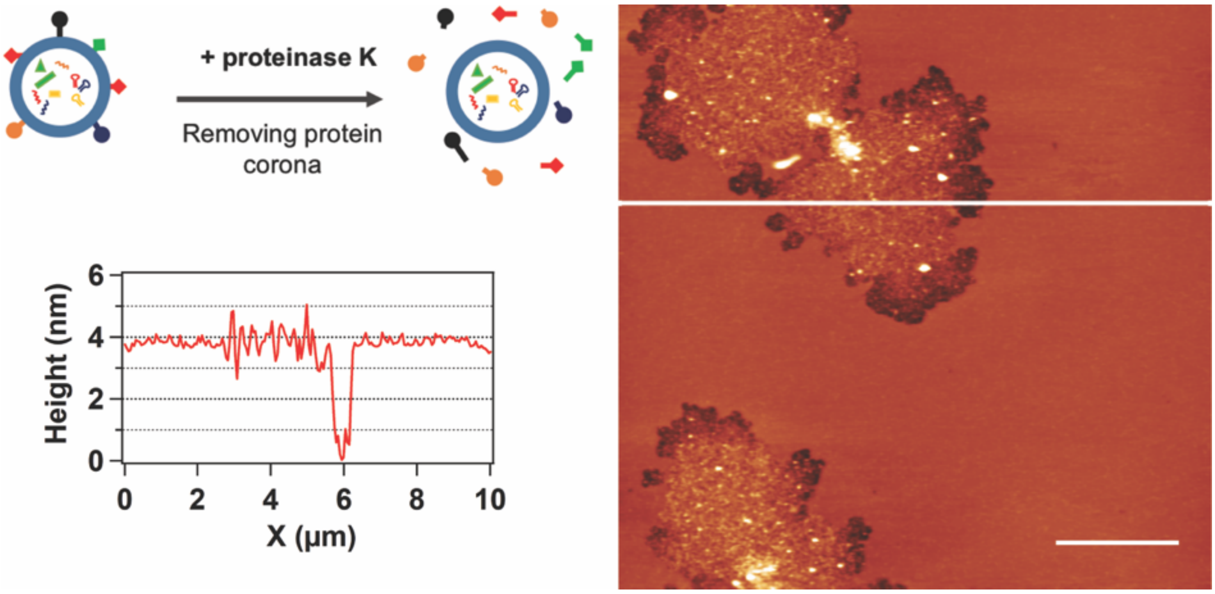


**Figure S4.** Atomic Force Microscopy topographic image and corresponding line profiles of DOPC SLB after addition of EVs previously treated with proteinase k, following the protocol reported by Skliar et al. (Ref. 42 Main manuscript)

| | **SLD heads in ($E^{-6}$ Å$^{-2}$)** | **Thick heads in (Å)** | **SLD chains ($E^{-6}$ Å$^{-2}$)** | **Thick chains (Å)** | **SDL heads out ($E^{-6}$ Å$^{-2}$)** | **Thick heads out (Å)** |
|---|---|---|---|---|---|---|
| **DMPC** | 1.7 | 6.2 | 6.4 | 30 | 1.7 | 8.5 |
| **DMPC+ EVs LC** | 1.9 | 6 | 6.3 | 28 | 1.9 | 12 |
| **DMPC+ EVs MC** | 1.8 | 5 | 6.3 | 25 | 2 | 13 |
| **DMPC+ EVs HC** | 1.9 | 5 | 6.5 | 23 | 2.1 | 14 |

**Table S3.** Best fit parameters for SANS scattering patterns for bulk d54-DMPC SUVs before and after the interaction with EVs at low, medium and high concentration

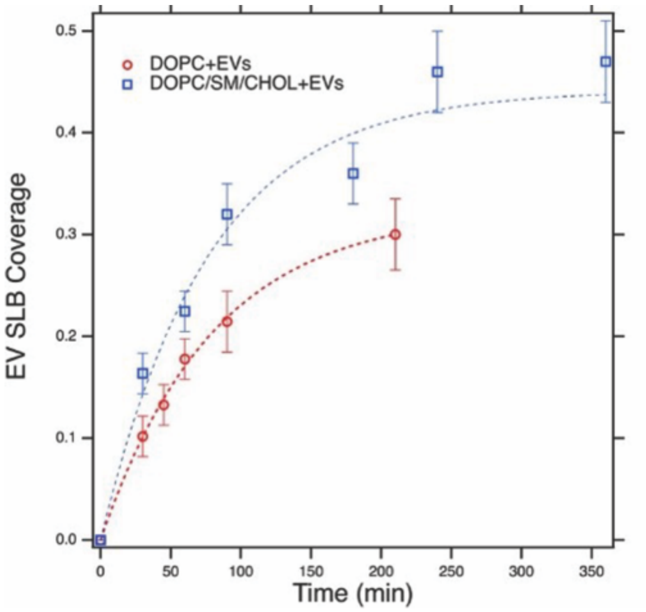


**Figure S5.** Evolution in time of the EV SLB coverage with respect to the total area of the artificial SLB for DOPC (red dots) and DOPC/SM/Chol (blue squares) systems. The dashed lines are fitting curves with a 1st order Langmuir model, $\Theta=A*(1-e^{-t/\tau})$, where $\Theta$ is the coverage, A is the maximum coverage and $\tau$ is the Langmuir adsorption model time constant. Fitting results for DOPC: A=0.32+/-0.05, $\tau$=80+/-20 minutes. Fitting results for DOPC/SM/Chol: A=0.44+/-0.03, $\tau$=77+/-11 minutes.
